# Systematic profiling of full-length immunoglobulin and T-cell receptor repertoire diversity in rhesus macaque through long read transcriptome sequencing

**DOI:** 10.1101/782938

**Authors:** Hayden N. Brochu, Elizabeth Tseng, Elise Smith, Matthew J. Thomas, Aiden Jones, Kayleigh Diveley, Lynn Law, Scott G. Hansen, Louis J. Picker, Michael Gale, Xinxia Peng

**Affiliations:** Department of Molecular Biomedical Sciences, North Carolina State University College of Veterinary Medicine, Raleigh, NC 27607, USA; Bioinformatics Graduate Program, North Carolina State University, Raleigh, NC 27695, USA; Pacific Biosciences, Menlo Park, CA 94025, USA; Department of Immunology, University of Washington, Seattle, WA, USA; Center for Innate Immunity and Immune Diseases, University of Washington, WA, USA; Genetics Graduate Program, North Carolina State University, Raleigh, NC 27695; Vaccine and Gene Therapy Institute, Oregon Health & Science University, Beaverton, OR 97006, USA; Washington National Primate Research Center, University of Washington, Seattle, WA, USA; Bioinformatics Research Center, North Carolina State University, Raleigh, NC 27695, USA

## Abstract

The diversity of immunoglobulin (Ig) and T-cell receptor (TCR) repertoires is a focal point of immunological studies. Rhesus macaques are key for modeling human immune responses, placing critical importance on the accurate annotation and quantification of their Ig and TCR repertoires. However, due to incomplete reference resources, the coverage and accuracy of the traditional targeted amplification strategies for profiling rhesus Ig and TCR repertoires are largely unknown. Here, using long read sequencing, we sequenced four Indian-origin rhesus macaque tissues and obtained high quality, full-length sequences for over 6,000 unique Ig and TCR transcripts, without the need for sequence assembly. We constructed the first complete reference set for the constant regions of all known isotypes and chain types of rhesus Ig and TCR repertoires. We show that sequence diversity exists across the entire variable regions of rhesus Ig and TCR transcripts. Consequently, existing strategies using targeted amplification of rearranged variable regions comprised of V(D)J gene segments miss a significant fraction (27% to 53% and 42% to 49%) of rhesus Ig/TCR diversity. To overcome these limitations, we designed new rhesus-specific assays that remove the need for primers conventionally targeting variable regions and allow single cell-level Ig and TCR repertoire analysis. Our improved approach will enable future studies to fully capture rhesus Ig and TCR repertoire diversity and is applicable for improving annotations in any model organism.

## Introduction

Rhesus macaque (*Macaca mulatta*) is one of the most commonly used and best-studied non-human primate (NHP) animal models. NHPs are key to studying human biology and human diseases due to their close phylogenetic relationship and similar physiology to humans [1–4]. They are frequently used for vaccine development [5] and to model infection with human pathogens, such as Mycobacterium tuberculosis [6, 7], HIV [8, 9], influenza A virus [10], and Zika virus [11], among others [12, 13]. Developing complete and accurate NHP genomic resources, especially for the immune system, is imperative for efficient translational interpretation [14, 15].

Critical for mounting adaptive immune responses, immunoglobulin (Ig) and T-cell receptor (TCR) repertoires house an enormous amount of diversity responsible for recognizing a near limitless array of antigens presented through environmental exposures. Ig and TCR have two domains: a constant region and a variable region. The repertoire diversity is mostly present in the variable region domain, which is comprised of a Variable (V), Joining (J), and in some cases a Diversity (D) gene segment (Fig. 1). Duplicated copies of these gene segments cluster into several large loci in the genome. As a result, correct assembly of their respective loci with highly repetitive sequences is a major technical challenge, one that has yet to be fully resolved in rhesus macaque despite significant improvements [16] and the recent release of a new genome assembly, rheMac10 (GCA_003339765.3). As demonstrated by a recent vaccine-related study in rhesus macaque [17], correct assembly of these complex regions requires much longer  databases (e.g. the international ImMunoGeneTics information system or IMGT) for these diverse sequences also remain fairly meager relative to their human counterparts, although there are new tools developed to address these gaps [18]. Since the design of available rhesus-specific assays for profiling Ig and TCR diversity has heavily relied on these limited rhesus reference resources, the coverage and the accuracy of these assays still require unbiased assessment and potentially improved approaches might be necessary. We begin by briefly summarizing the complexity of these immune repertoires in general and how this complexity has constrained the development of resources and assays for rhesus macaque.

**Fig 1.**
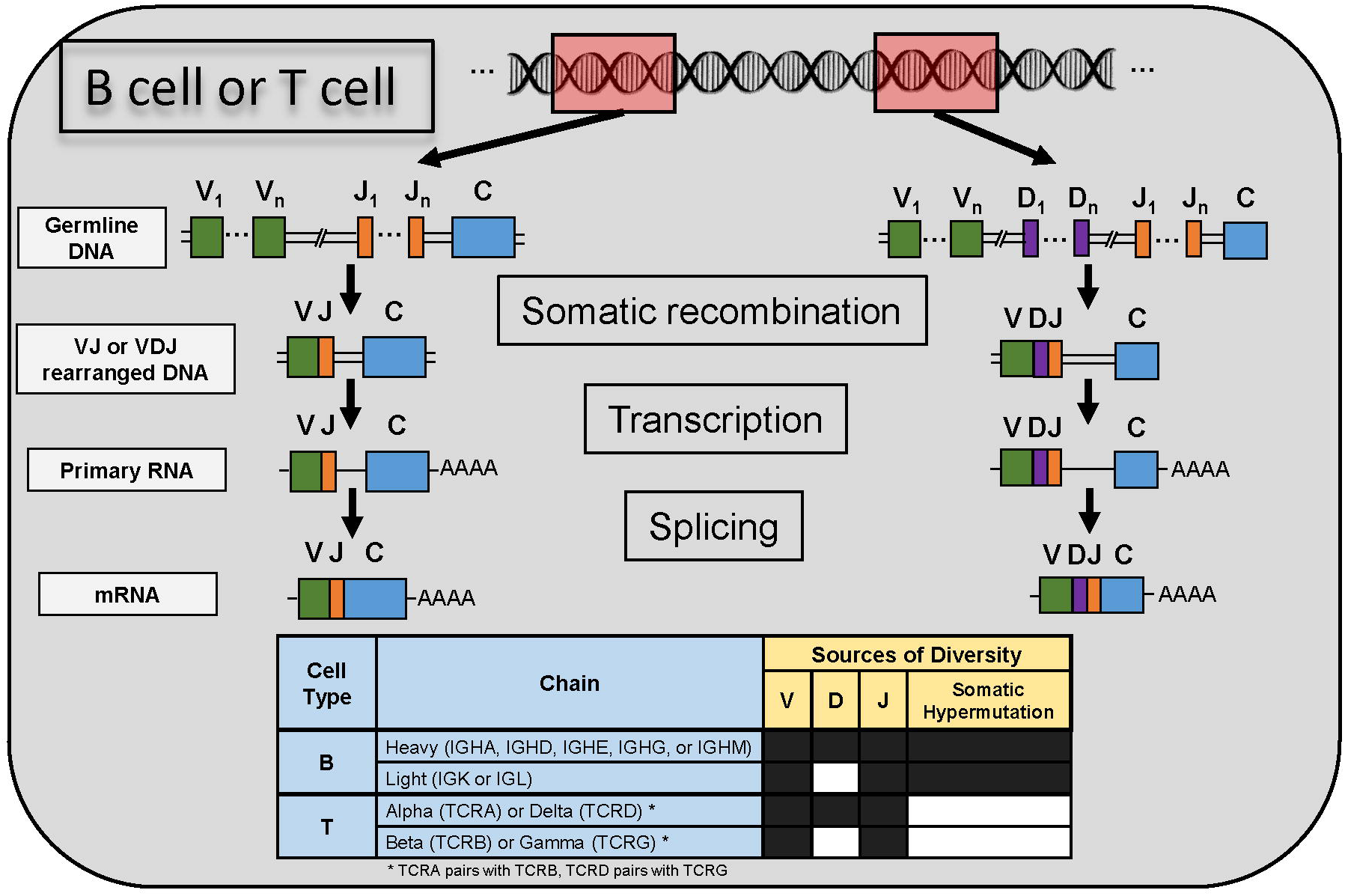
Overview of how immunoglobulin (Ig) and T cell receptors (TCR) form, beginning with germline DNA. Within a single cell (B or T), two chains are generated separately through either VJ or VDJ rearrangement (somatic recombination) at the DNA level. These molecules are then transcribed and spliced to form mRNA molecules that include a constant region (C) domain. The constant region is what determines the specific chain type shown in the table beneath the graphic. For example, in a B cell, Ig consists of a heavy chain and a light chain. Depending on its constant region, the heavy chain is further classified as alpha (IGHA), delta (IGHD), epsilon (IGHE), gamma (IGHG), or mu (IGHM), while the light chain is either kappa (IGK) or lambda (IGL). The sources of diversity are analogous across B and T cells, with the exception of the former undergoing somatic hypermutation, a process that introduces random mutations in the variable (VJ or VDJ) region.

In individual B and T cells, different gene segments of both variable and constant regions are combined at the DNA level to encode distinctive functional Ig and TCR genes through a process known as somatic recombination (Fig. 1) (reviewed in [19–21]). Somatic recombination accounts for the majority of the diversity within the variable region domain, with the number of unique Ig and TCR variable region domains estimated to be on the order of 10^13^ and 10^18^, respectively [22]. To further increase the repertoire diversity, within each cell a fully assembled Ig molecule is composed of a unique pairing of a heavy chain (IGH) and a light chain: either kappa (IGK) or lambda (IGL) (Fig. 1). Similarly, a TCR is composed of the pairing of an alpha (TCRA) and beta (TCRB) chain or, less commonly, a delta (TCRD) and gamma (TCRG) chain (Fig. 1). The process of somatic hypermutation (SHM) further expands the diversity of Ig molecules, where mutations are introduced to the variable region domain of recombined DNA at rates roughly one million times higher than background rates observed in other genes [23].

The constant region domain determines various chain types of Ig and TCR [22, 24]. Within each chain type, these variable gene segments are classified into families (e.g. Ig heavy chain variable gene family 4 or IGHV4). Each gene family includes one or more unique gene segments (e.g. IGHV4-1) that share high degrees of similarity. In B cells, the process by which the variable region domain is juxtaposed from the originally utilized heavy (H) chain constant region gene (usually IGHM) to a different constant region domain is referred to as class-switch recombination (CSR), a process essential to Blymphocyte differentiation. In addition to conferring the non-antigen binding functions of antibody molecules, the heavy chain constant region of secreted Ig differs from Ig bound to the surface of B cells (the B cell receptor for antigen) based on alternative splicing at the 3’ end.

The genetic diversity of the Ig and TCR loci present a unique challenge for accurate measurement. Traditionally, these immune repertoires are targeted for amplification either by multiplex polymerase chain reaction (MPCR) [25], RNA capture [26], or 5’ rapid amplification of cDNA ends (5’ RACE) [27]. For rhesus macaque, many Ig repertoire sequencing efforts use a MPCR approach [28, 29], while some employ 5’ RACE [30]. Typically, such PCR-based approaches are designed for individually sorted B or T cells [28] and facilitate cloning efforts [31]. A more recent rhesus-specific MPCR design aimed to expand coverage of the Ig repertoire [32]. There were also attempts to improve rhesus V and J germline gene annotation using 5’ RACE sequencing (RACE-seq) [18, 33]. The recently developed IgDiscover tool [18] now makes it possible to leverage germline databases of related species to improve those of model organisms, for example using human germline databases to study Indian- and Chinese-origin rhesus macaques. Authors from the same lab also developed a strategy for targeting the 5’ UTR of V genes, often conserved among these gene clusters, thus reducing the number of primers needed to target the variable region in human [34]. However, rhesus MPCR amplification systems are inherently biased to the V and J gene segments they target because the primers are designed based on the consensus of a limited number of reference sequences [35].

RACE-seq has the advantage of only targeting the constant region, where primer design is significantly easier given the low sequence variability. 5’ RACE in addition to MPCR have also been modified to incorporate unique molecular identifiers (UMIs) for repertoire sequencing [36, 37], mitigating issues arising from PCR bias [38]. However, such protocols are optimized for applications in human and mouse [39] and have not yet been applied for Ig/TCR analysis in rhesus macaques. Moreover, even with MPCR or 5’ RACE it is still difficult to capture complete Ig mRNA transcripts with the commonly available sequencing instruments like the 2 x 250 bp HiSeq system and the 2 x 300 bp MiSeq system, in part due to the large length variability of recombined molecules [40] and the potential for non-canonical recombination events [41]. Library preparation techniques have recently been improved to support this capability but only in human [34]. Despite these technical advances, it is still unclear how well these strategies perform in rhesus macaque, where the relatively limited resources preclude similar benchmarking efforts previously done for human Ig repertoires [42] and TCR repertoires [43].

The technical robustness and unique advantages of scRNA-seq [44, 45] have revolutionized the analysis of human immune systems [46–50], resulting in an increasing number of studies for single cell-based Ig and TCR repertoire sequencing in human [51–54]. Single cell protocols target repertoire sequences similarly to RACE-seq by only targeting the constant regions of sequences and by incorporating UMIs. Interrogation of Ig/TCR repertoire and B/T cell transcriptomes in the same single cells has provided novel insights in human, such as evaluating T cell clones within tumors [51], modeling of epitope specificities [52, 53], and assessment of host-microbiome immune homeostasis [54]. However, to the best of our knowledge, for NHP models including rhesus macaque, equivalent assays for single cell Ig and TCR repertoire analysis are still not available.

In this study, we sought to address the technical limitations of existing immune repertoire sequencing protocols in rhesus macaque. To circumvent the highly difficult task of correctly assembling Ig/TCR transcripts from short sequencing reads, we performed long read transcriptome sequencing of rhesus tissue samples. We constructed the first complete reference set for the constant regions of all known isotypes and chain types of rhesus Ig and TCR repertoires from high-quality, full-length Ig and TCR transcript sequences. Our resource provides the multiple rhesus Ig and TCR constant regions still missing in the current IMGT database, avoiding complications due to incomplete assemblies of their respective genomic loci. Using this unbiased collection of full-length Ig and TCR transcript sequences, we were able to examine the diversities across the entire variable regions of rhesus Ig and TCR transcripts and the coverage of available assays for profiling rhesus Ig and TCR repertoires. Further, we designed new rhesus-specific assays that allow much broader rhesus Ig and TCR repertoire analysis both at the single cell level and using 5’ RACE. These results furthermore demonstrate that immunoprofiling resources and assays can be developed for other organisms similarly without complete genomic assemblies of immune loci.

## Materials and Methods

### Tissue sample preparation and transcriptome sequencing

To construct cDNA libraries for PacBio SMRT sequencing, we used previously isolated Indian-origin rhesus macaque RNA from four tissue types: lymph node (LN), peripheral blood mononuclear cells (PBMC), whole blood (WB), and rectal biopsy (RB). To maximize the coverage of transcript diversity, we pooled RNA from multiple Indian-origin rhesus macaques. The pooled WB RNA was from six healthy, uninfected macaques, while the pooled LN, PBMC, and RB RNA were from 18 macaques infected with SIVmac251. The LN RNA came from a combination of peripheral and proximal mesenteric lymph nodes. Using a Clontech SMARTer kit, cDNA was produced from each pooled RNA sample followed by PCR amplification. The amplified cDNA samples were size selected using BluePippin (Sage Science) into four size fractions: 1-2 kb, 2-3 kb, 3-6 kb, >6 kb. Libraries were prepared using the SMRTbell template prep kit 1.0 and sequenced on the PacBio RS II platform with P6-C4 chemistry and 4-hour movie times at the University of Washington PacBio Sequencing Services site.

### Data processing

Raw PacBio sequencing data was first run through the CCS protocol in SMRT Analysis 2.3.0 to generate Circular Consensus Sequence (CCS) reads. Each CCS read is the circular consensus sequence of a single molecule. CCS reads were classified as non-full-length and full-length, where the latter has all of the following detected: polyA tail, 5’ cDNA primer, and 3’ cDNA primer. full-length and non-full-length CCS sequences were aligned to immunoglobulin (Ig) and T cell receptor (TCR) databases using IGBLAST v1.8.0 [55] and IMGT annotation [56]. Separate searches were carried out using either human or rhesus macaque germline V, D, and J sequences where available (both for Ig, only human for TCR). Full-length Ig hits were kept for downstream analyses when they were deemed functional by IgBLAST output, meaning they lacked premature stop codons and were in-frame. When Ig hits had both rhesus and human database hits, the rhesus annotation was used in all subsequent analyses.

### Generation of consensus constant region sequences

To validate the chain types reported by IgBLAST and gather additional information about rhesus Ig/TCR isotypes and their subclasses, we further processed full-length Ig and TCR constant region sequences as below. Using the J region reported by IgBLAST, we extracted the constant region sequence for each full-length sequence. Next, we clustered extracted constant region sequences using CD-HIT [57] with the following parameters: -c 0.97 -G 0 -aL 0.95 -AL 100 -aS 0.99 -AS 30 (https://github.com/Magdoll/cDNA_Cupcake/wiki/Tutorial:-Collapse-redundant-isoforms-without-genome). Ig and TCR sequences were clustered separately. We only kept clusters with at least three constituent sequences for downstream consensus sequence analysis. For those clusters with at least ten sequences we also performed a second round of more stringent clustering using a 99% identity cutoff (-c 0.99 with CD-HIT) to separate quality sequences from those with multiple indels and to identify potential allotypes. From the second round of more stringent clustering, consensus sequences were generated using only the sequences from the largest remaining clusters, with the goal of removing low quality sequences that formed many small clusters. When this second round of clustering resulted in multiple large clusters of comparable size, a consensus sequence was generated for each cluster. We used the cons tool from the EMBOSS v6.6.0 suite [58] to generate consensus sequences, identifying and removing any remaining insertions at predominantly gapped locations. Protein sequences were derived from the generated consensus sequences using the EMBOSS transeq tool [58], where the frame with the longest protein sequence was kept. We verified their identities via global alignments of the coding regions of cDNA sequences to a database of IMGT human constant region nucleotide coding regions [56] using the usearch_global tool in the USEARCH suite [59], considering only the best alignment. We aligned consensus sequences belonging to the same chain type (and isotype if applicable) using Clustal Omega [60], and removed any remaining redundancies in the consensus sequences after manual visualization and curation in UGENE [61].

Next, we used these databases to systematically classify putative Ig and TCR sequences using all CCS reads from the complete Iso-Seq dataset as input. CCS reads were separately globally aligned to the custom databases of consensus Ig and TCR constant region sequences using the usearch_global tool [59] with a 90% identity threshold to ascertain chain type and isotype/subclass where appropriate. Sequences without large gaps (>10bp) in their alignment were kept in the final assignment of Ig and TCR sequences, as it was unclear if those sequences with large gaps were rare alternatively spliced transcripts or the result of sequencing errors. We then compared the CCS reads confirmed as Ig/TCR by our custom databases with the CCS reads identified by the initial IgBLAST search. Based on these comparisons, the final set of Ig transcripts were filtered to only include CCS reads that were confirmed by the custom database and also IgBLAST hits deemed functional by rhesus IMGT annotation (Supplementary Fig. 1B). Further, the final set of TCR transcripts only required CCS reads to be confirmed by the custom database (Supplementary Fig. 1A), as it was unclear if IMGT functional annotation for human would be informative for rhesus-derived transcripts. We quantified the rates of insertions and deletions in the constant regions of confirmed Ig and TCR sequences using the CIGAR strings in the global alignments of CCS reads against this custom constant reference database by usearch_global [59]. Non Ig/TCR CCS reads of this complete Iso-Seq dataset are being analyzed in a separate study.

### Analysis of V-J usage and variable region diversity

The frequencies of V-J combinations were measured using the IMGT annotation reported by IGBLAST [55, 56] for all final Ig transcript sequences ascertained in the above analysis. Chi-square test for independence based on V-J combination frequencies was performed in R [62] and by requiring that at least 50% of elements in each row and column have at least 10 counts. The diversity of the variable region sequences was determined by first removing the constant region sequence determined by its alignment to the corresponding consensus constant region sequences. Separate multiple sequence alignments of the most common V-J regions were performed using Clustal Omega [60] and the consensus profiles were visualized in R with ggplot2 [62, 63].

### *In silico* PCR analysis

Common rhesus-specific MPCR assays for amplifying IGH, IGK, and IGL sequences [28, 29, 32] and TCRA and TCRB sequences [64, 65] were tested via *in silico* PCR analysis using the UCSC isPcr tool [66]. For those primer sets with a nested design [28, 29], only the inner primers were used for the analysis. The standard inner and outer reverse primers used for the 10x B cell and T cell V(D)J assays were also assessed via *in silico* PCR analysis [66] where a dummy adapter sequence was prepended to the 5’ end of sequences to enable analysis with the isPcr tool and only test reverse primer efficiencies. The efficiencies of rhesus-specific MPCR assays [28, 29, 32] in amplifying the variable region (forward primer) and constant region (reverse primer) were similarly tested by either appending and prepending a dummy adapter sequence, respectively. In all uses of the isPcr tool, default parameters were used, requiring a perfect match for the first 15 nucleotides on the 3’ end of the primer.

### Design of rhesus-specific primers and PCR validation

Ig and TCR constant region inner and outer reverse primers were designed using NCBI primerBLAST [67] with target Tms consistent with current 10x V(D)J assay forward primers. For those targeting multiple consensus sequences, the search space was constrained to consensus regions. To validate these 10x optimized reverse primers, primerBLAST [67] was used to design standard PCR assays each with a forward primer compatible with its respective 10x optimized reverse primer and target reference sequence. This was only done for custom primers in the V(D)J assays, while primers adopted from Sundling et al [28] were assumed to be efficacious. 100 nmole HPLC purified DNA oligonucleotides were ordered from IDT. Reverse transcription was performed using Qiagen QuantiTect Reverse Transcription Kit (Cat. No. 205313, Lot 157037104) following the manufacturer’s protocol with 1 ug rhesus macaque lymph node tissue RNA. The resulting cDNA was then amplified using Takara Titanium TaqPCR kit (Cat. No. 639210, Lot 1805465A) following the manufacturer’s protocol (https://www.takarabio.com/assets/documents/User%20Manual/PT3304-2.pdf), altering the recommended cycling conditions with 30 cycles and an annealing temperature and time of 50°C and 1 minute. 400 ng of each PCR reaction was then run on a 10% TBE polyacrylamide gel alongside Thermo Scientific 100 bp GeneRuler to confirm product size from primer pairs.

## Results

### Generation of a complete reference collection for rhesus macaque Ig and TCR constant regions

Using PacBio transcriptome sequencing (the Iso-Seq method), we obtained over 2.8 million Circular Consensus Sequence (CCS) reads from four different rhesus macaque tissues (Supplementary Table 1). Each CCS read is the circular consensus sequence of a single transcript molecule [68]. About 33% of these CCS reads were full-length, i.e. contained the 5’ cDNA primer, 3’ cDNA primer, and polyA tail (Supplementary Table 1). Using the available IMGT annotation of the variable regions of Ig (human and rhesus) and TCR (human) germline sequences [56] and IgBLAST [55], we recovered 13,118 Ig and 2,534 TCR putative transcript sequences, respectively accounting for 1.4% and 0.28% of the total number of full-length CCS reads (Supplementary Table 1). Only functional Ig transcripts (i.e. those in-frame and without premature stop codons according to IMGT germline annotation) were used in downstream analyses (5,666 / 13,118 or 43% of full-length Ig transcripts). We kept all full-length TCR transcripts at this step, as it was unclear if their functionalities could be correctly determined with the use of human germline TCR reference sequences.

We first sought to characterize the constant regions of these full-length Ig and TCR transcript sequences. Since constant regions are significantly less variable, this would allow us to compile a complete reference set and accurately classify the different chain types and isotypes. Using the V(D)J annotation provided by IMGT [56], we extracted constant regions from the full-length Ig and TCR transcript sequences. We then clustered and curated these sequences to generate complete cDNA and coding sequences (Materials and Methods). Complete cDNA sequences represent the entire constant region consensus (i.e. including the 3’ UTR), while the coding regions were generated at both the cDNA and protein level. We obtained the complete consensus sequences for the following chain types and isotypes with multiple references noted in parentheses: IGHA (5), IGHD, IGHE, IGHG1, IGHG2, IGHG3, IGHG4, IGHM (2), IGLC, IGKC, TCRAC, TCRBC1, TCRBC2, TCRDC, TCRGC1, and TCRGC2. We thus recovered complete reference sequences for all known secreted IGH isotypes, and additionally two IGHA and one IGHM membrane-bound reference sequences. We numbered membrane-bound rhesus Ig reference sequences to be consistent with their corresponding secreted forms.

We identified three unique IGHA consensus sequences: two in both secreted and bound form and one in only secreted form. Much of the variation among these sequences was confined to the hinge region, which has previously been reported to be highly heterogeneous among rhesus macaques [69]. In fact, we recovered three of the eight unique hinge regions previously identified by [70] (data not shown). We numbered these unique IGHA sequences based on the relative abundances we later determined within our samples (i.e. IGHA*01 was the most abundant), following conventional naming for allelic variants.

We also identified all known subclasses of IGHG, TCRBC, and TCRGC. We aligned the four IGHG cDNA coding sequences we recovered to the available rhesus IGHG gDNA in IMGT to determine their subclasses. Each cDNA coding sequence reference corresponded to one of the four IGHG subclasses with high sequence identities (98.6% to 99.9%). To properly classify the TCRBC subclasses, we aligned the two TCRBC protein sequences to the two reference ORFs available in IMGT, yielding perfect matches (data not shown). We leveraged the available human ORFs in IMGT to determine subclasses of TCRGC, as reference sequences were not available for rhesus macaque. Using a multiple sequence alignment, we assigned orthologous subclass naming to the rhesus references based on the human reference with the highest structural similarity (data not shown).

Interestingly, 39% (979 of 2,534) of putative TCR transcript sequences identified by the initial IgBLAST search were in fact Ig transcripts, evidenced by recapitulation of several Ig consensus sequences among the set of TCR clusters and by successful alignment of such sequences to the set of Ig reference sequences (Supplementary Fig. 1A). While this did not prevent our complete recovery of TCR consensus sequences, it suggests that commonly used human reference based IgBLAST searches could be inaccurate for rhesus TCR repertoire analysis.

### Classification of rhesus Ig and TCR transcript sequences

Next, we used these consensus sequences of rhesus constant regions to classify the full-length rhesus Ig and TCR transcript sequences. When we aligned the constant regions of full-length functional Ig transcripts to this newly constructed database of rhesus Ig cDNA consensus sequences, we obtained successful assignments for 96% (5,415 / 5,666) of these full-length Ig transcript sequences. Those transcript sequences that failed to align tended to be other molecules from the immunoglobulin superfamily that contain V-set domains (i.e. a variable region), representing false positives from our initial IgBLAST screen (data not shown). Among those aligned, a small fraction (1%) of sequences had large structural differences, indicating infrequent alternative splicing events (data not shown). To simplify the downstream analysis, we removed these transcripts from further processing, yielding a final set of 5,384 full-length Ig transcript sequences (Table 1). Among four tissue samples sequenced here, the rectal biopsy and lymph node had the largest number of unique Ig transcript sequences overall. Secreted IGHA and IGHG transcripts had the highest relative abundance and were most prevalent in the rectal biopsy and lymph node samples. IGK and IGL sequences were significantly less abundant than IGH isotypes in general, but they were still detected across each tissue.

**Table 1.** Number of unique full-length rhesus IG transcript sequences identified in each tissue. Counts for secreted heavy chain Ig sequences (IGH A, D, E, G1-4, and M) are shown with membrane-bound counts in parenthesis where applicable. IGHA is further classified by allotype, conventionally denoted by an asterisk followed by a number.

|  | Rectal Biopsy | Whole Blood | PBMC | Lymph Node | Total |
| --- | --- | --- | --- | --- | --- |
| IGHA*01 | 1411 (8) | 67 (0) | 21 (2) | 224 (0) | 1723 (10) |
| IGHA*02 | 705 (4) | 7 (0) | 21 (1) | 275 (6) | 1008 (11) |
| IGHA*03 | 101 | 0 | 5 | 9 | 115 |
| IGHD | 0 | 12 | 4 | 3 | 19 |
| IGHE | 1 | 1 | 0 | 4 | 6 |
| IGHG1 | 78 | 21 | 21 | 1030 | 1150 |
| IGHG2 | 9 | 9 | 4 | 311 | 333 |
| IGHG3 | 0 | 1 | 0 | 30 | 31 |
| IGHG4 | 1 | 0 | 0 | 13 | 14 |
| IGHM | 129 (3) | 20 (21) | 18 (14) | 169 (8) | 336 (46) |
| IGK | 97 | 30 | 12 | 155 | 294 |
| IGL | 100 | 30 | 18 | 140 | 288 |
| Total | 2632 (15) | 198 (21) | 124 (17) | 2363 (14) | 5317 (67) |

Similarly, we identified rhesus TCR transcript sequences by aligning raw full-length CCS reads directly to this custom collection of rhesus TCR cDNA consensus sequences. This precluded any bias against rhesus TCR transcript sequences that contained V, D, and J genes with low similarity to those in the human IMGT reference. In total, we assigned 741 full-length CCS reads as rhesus TCR transcript sequences (Table 2), 50 of which (7%) we did not originally recover using the human IMGT reference database (Supplementary Fig. 1A). This suggests there was a significant divergence between human and rhesus TCR germline V(D)J genes, indicating that a species-specific germline database is necessary for complete repertoire recovery. We detected the fewest TCR transcript sequences in the rectal biopsy sample, while the whole blood, PBMC, and lymph node samples each had larger recoveries of all TCR isotypes (Table 2). TCRA was the most abundant in each tissue, representing 61% of all sequences recovered (455 / 741). TCRG2 had the second highest abundance with its strongest representation in PBMCs (Table 2).

**Table 2.** Number of unique full-length rhesus TCR transcript sequences identified in each tissue.

|  | Rectal Biopsy | Whole Blood | PBMC | Lymph Node | Total |
| --- | --- | --- | --- | --- | --- |
| TCRA | 14 | 71 | 103 | 267 | 455 |
| TCRB1 | 1 | 9 | 11 | 26 | 47 |
| TCRB2 | 0 | 13 | 16 | 44 | 73 |
| TCRD | 1 | 1 | 27 | 10 | 39 |
| TCRG1 | 2 | 3 | 6 | 5 | 16 |
| TCRG2 | 3 | 11 | 75 | 22 | 111 |
| Total | 21 | 108 | 238 | 374 | 741 |

We then assessed the sequence quality of these Ig and TCR transcripts, by quantifying insertion and deletion events within the alignments of their constant region sequences and the corresponding reference cDNA sequences. We elected not to evaluate the rate of substitutions, as they are difficult to discern from allelic variation given our sequencing depth and are generally less common than insertions and deletions in CCS reads [33]. The rate of insertions in Ig constant region sequences was very low, with a mean of 0.11% and median of 0.06%. The mean and median deletion rates were slightly lower at 0.04% and 0%, respectively. Error rates within the TCR constant regions were comparable to those of Ig constant regions, with mean insertion and deletion rates of 0.15% and 0.07% and median rates of 0.07% and 0%, respectively. These low error rates reflected the high quality of the final set of full-length rhesus Ig and TCR transcript sequences we obtained in this study.

We found the recovery of TCR transcripts to be significantly more accurate and marginally more sensitive when using our custom TCR constant region database for identification instead of the traditional IgBLAST approach with human germline annotation (Supplementary Fig. 1A). We thus elected to align CCS reads directly to our custom Ig constant region database to make a similar comparison. We recovered 11,451 Ig transcripts using this strategy; of these, 5,384 (47%) were functional IgBLAST hits (see Table 1), 5,858 (51%) were non-functional IgBLAST hits, and 209 (2%) were not detected by IgBLAST (Supplementary Fig. 1B). B cells that contain non-functional receptor sequences are known to apoptose early in their development; therefore, the proportion of non-functional IgBLAST hits observed here may be a reflection of the B cell population captured with these full-length transcripts. Many of the full-length Ig transcripts that eluded IgBLAST detection had enough sequence upstream of the constant region to harbor a complete variable region (127 or 1% overall) (Supplementary Fig. 1B), indicating that some variable region genes may be significantly diverged from those annotated in IMGT.

### Usage of known V, D, and J genes in Ig transcripts indicates broad coverage of rhesus Ig repertoire

To assess our overall coverage of the rhesus Ig repertoire, we examined the usage of individual V, D, and J genes among these full-length rhesus Ig sequences based on the alignment of their variable regions to rhesus Ig germline annotations in IMGT [56]. We detected all known gene families of rhesus IGHV, IGHD, and IGHJ genes among the four tissue samples (Fig. 2). The majority of IGHV and IGHJ genes detected were from the IGHV4 and IGHJ4 families, respectively, while there was broad coverage of IGHD gene families. Among light chain gene families, IGKV1 and 2 as well as IGLV1, 2, and 3 were the most frequent. IGKJ families were relatively less skewed in frequency, though IGLJ1 was in slightly higher frequency than other IGLJ families. The only known gene families we did not detect in these tissue samples were IGKV5, IGLJ4, and IGLJ5, likely due to the overall lower abundance of light chain Ig sequences we observed (Table 1). Interestingly, relative frequencies of these different gene families were highly consistent across the four tissues analyzed here (Supplementary Fig. 2), despite drastically different sampling depth (Fig. 2). Only light chain gene families within PBMCs and whole blood deviated from this trend. Since we had significantly less detection of IGK and IGL sequences in these tissues (Table 1), their observed deviations require further investigation.

**Fig 2.**
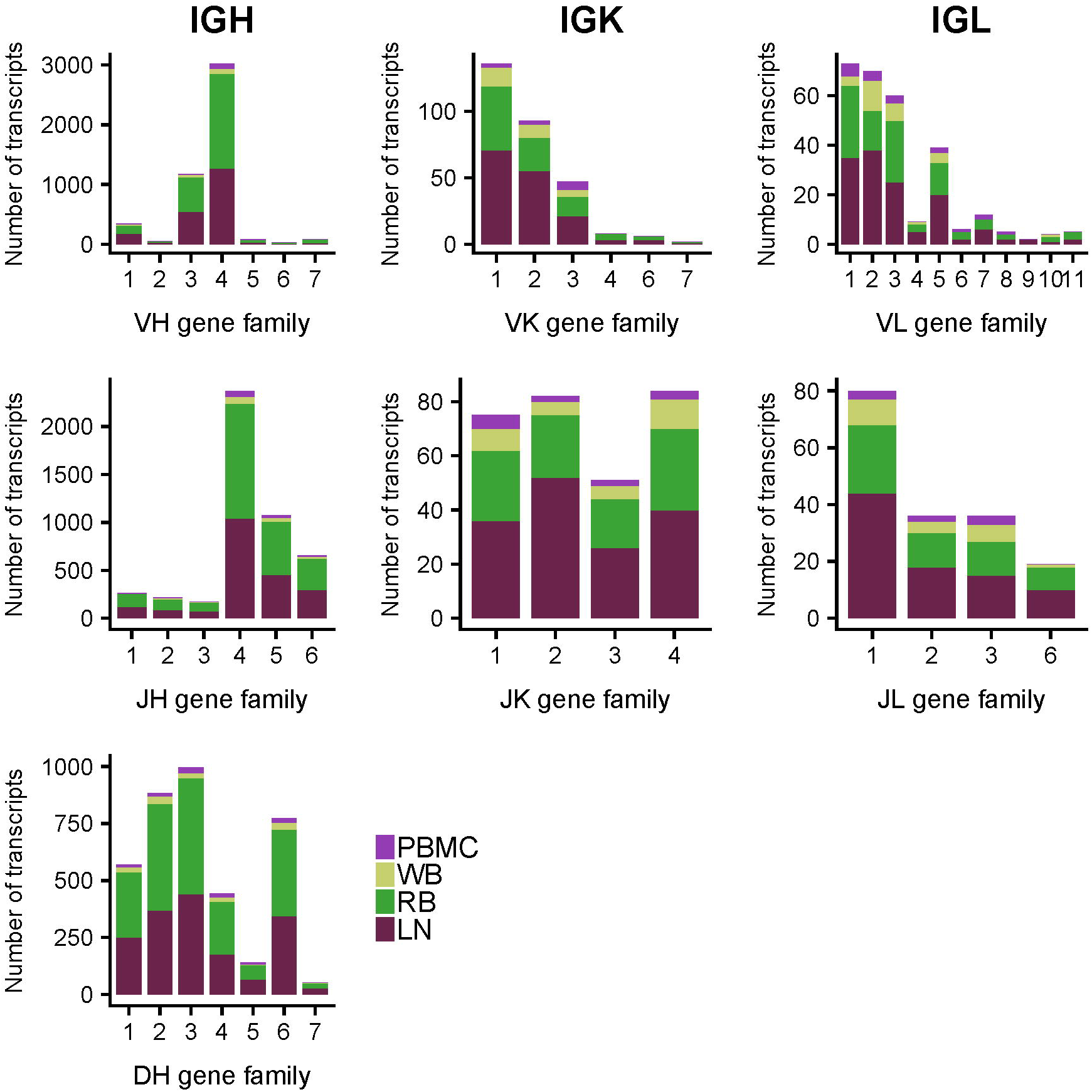
Stacked bar plot of absolute transcript abundance of V, D, and J genes for each Ig chain type, where colors of the bar indicate the tissue source of the gene. Variable gene abundance was computed using full-length Ig transcripts reported in Table 1. Each individual plot shows the various known gene families of a particular gene segment type (e.g. VH), where gene families are clusters of highly similar gene segments within that gene segment type.

Given the sufficient depth of coverage observed for IGH V, D, and J gene families, we calculated the combination frequencies of these V and J genes. Consistent with V and J gene frequencies observed in Fig. 2, the majority of V-J combination events contained IGHV4-2 and/or IGHJ4 (Fig. 3). Using the most abundant V and J genes, we tested the independence of V-J recombination events. We observed a significant nonrandom distribution of V-J recombination frequencies (chi square = 77.6, df = 15, p = 1.9e-10). Furthermore, we discovered a strong positive correlation between the V-J recombination rates in the lymph node and rectal biopsy samples (Pearson correlation = 0.86). Since these two samples had highly different compositions of IGH isotypes (Table 1), we also reasoned that there might be no discernable association between the variable regions and constant regions in general.

**Fig 3.**
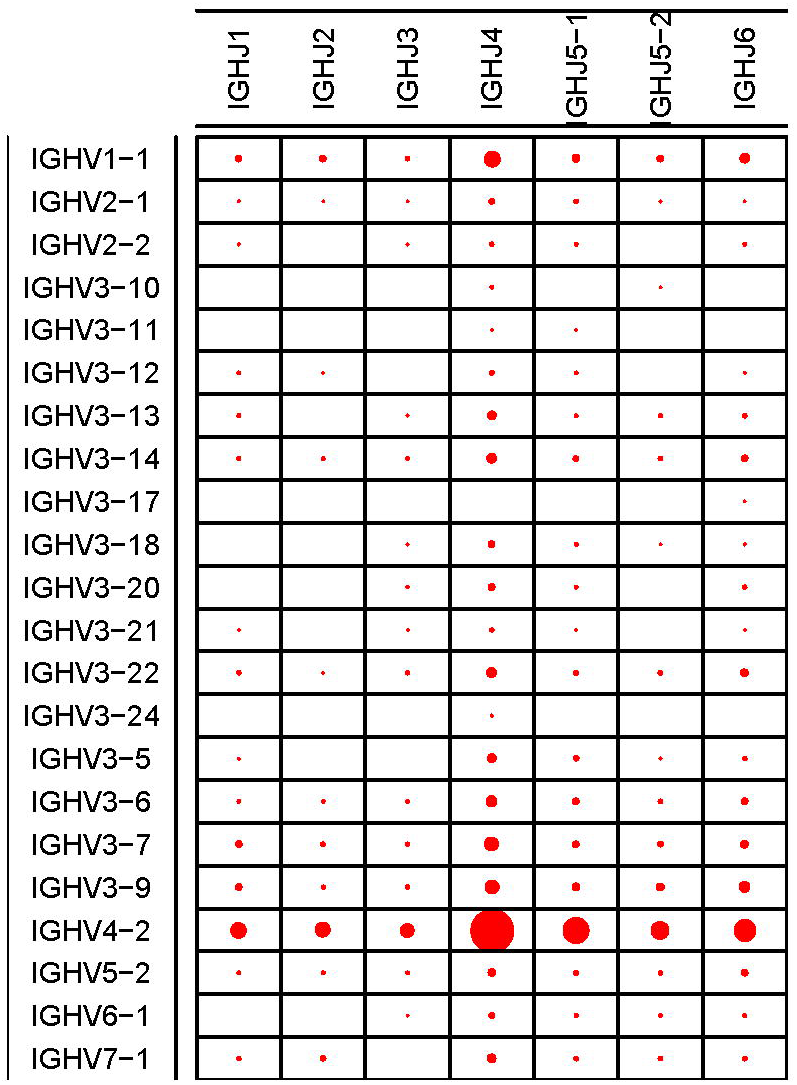
Dot plot of various IGHV and IGHJ combinations. The sizes of dots are proportional to the relative frequencies among the full-length Ig transcript sequences. The gene segments are conventionally named according to gene family (e.g. IGHV1) followed by the specific gene segment number. Gene segments simply have the family gene name (e.g. IGHJ1) when there is only one known gene segment in the family.

### Sequence diversity existing across the entire variable region shows MPCR amplification of rhesus immune repertoires is inherently biased

The collection of unbiased full-length Ig and TCR transcript sequences also provided a unique opportunity for exploring the sequence diversity across the entire variable region, the target of current repertoire amplification efforts. We assessed the efficiencies of three commonly used rhesus-specific Ig MPCR strategies [28, 29, 32], by examining the sites targeted by their primers in the context of their respective Ig V genes. We selected the three most abundant IGHV genes represented among the unique Ig transcripts: IGHV4-2 (1,383 sequences), IGHV1-1 (163 sequences), and IGHV3-9 (128 sequences), and generated consensus profiles for each. We discovered that the rhesus primers designed to target these genes all locate in regions rich in sequence diversity (Fig. 4). For example, Sundling et al [28] targeted well-conserved regions of IGHV4-1 and IGHV1-1 (Fig. 4A-B), yet appeared to target a region in IGHV3-9 that had a low percentage of sequences sharing the same consensus nucleotides (Fig. 4C). The low consensus observed in all three profiles at the 5’ end of the cDNA sequence (left side of profiles) was likely a result of the lack of guaranteed 5’ capture among CCS reads. However, it was confined to regions upstream of the primer target sites (Fig. 4A-C) and thus did not affect our assessment of available primers.

**Fig 4.**
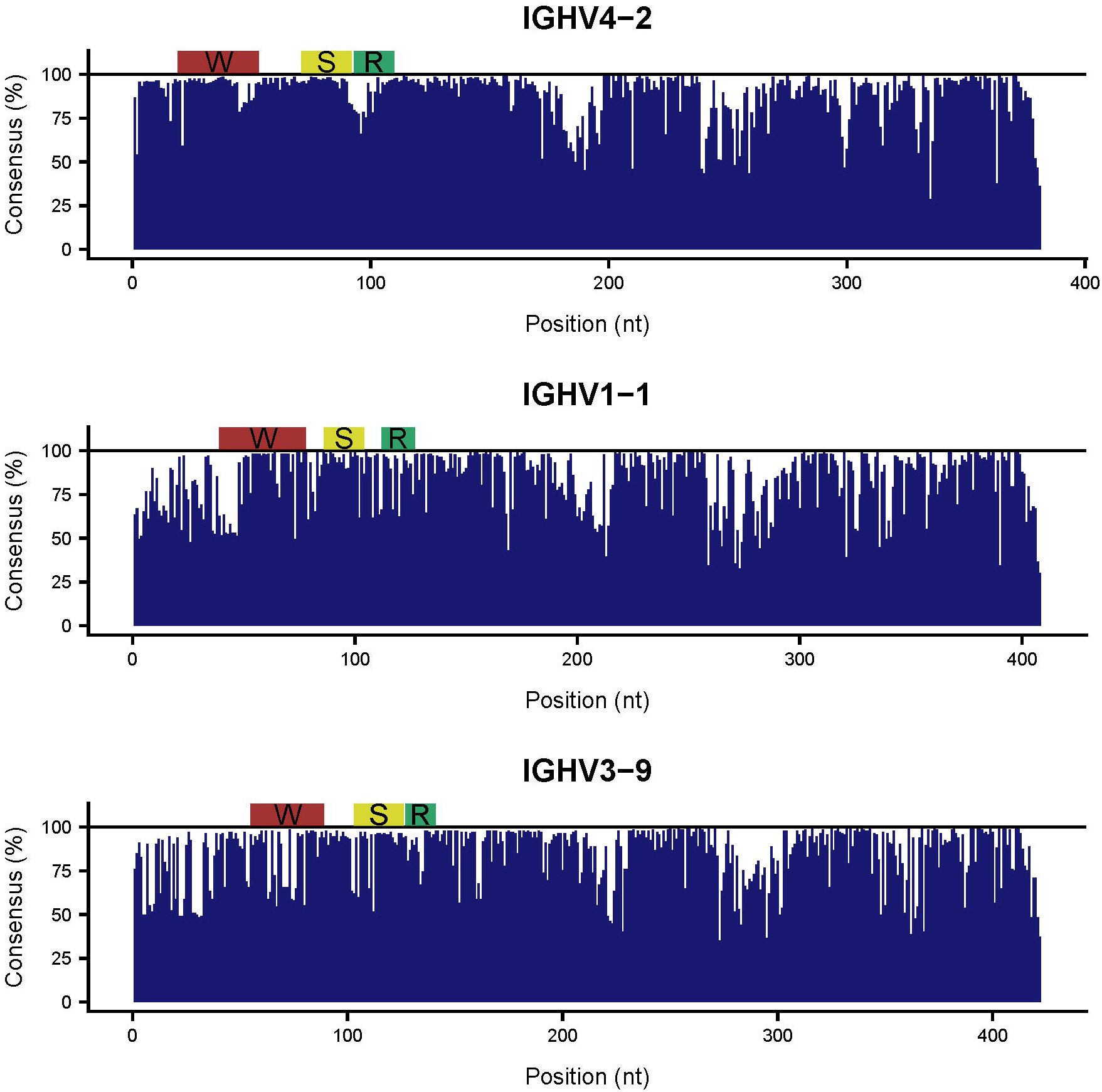
Consensus profiles of the most abundant IGHV genes detected. Panels **A-C** respectively show the percentage of nucleotides at each position that match the consensus (most abundant) nucleotide for IGHV4-2, IGHV1-1, and IGHV3-9. The IGHV4-2 consensus was constructed from 1,383 sequences, while IGHV1-1 was from 163 sequences and IGHV3-9 was from 128 sequences. The regions targeted by primers from (S)undling et al [28], (W)iehe et al [29], and (R)osenfeld et al [32] respectively have either a yellow, red, or green bar above.

To illustrate the limitation that the observed diversity could reduce the overall coverage of these MPCR approaches, we evaluated these primer sets via *in silico* PCR analysis with the full-length Ig transcript sequences as the source of potential template transcripts (Fig. 5). Similarly, we also leveraged known rhesus-specific MPCR strategies for TCR [64, 65] in this analysis (Fig. 5). We first tested only the forward (V gene) primers from these Ig and TCR primer sets [28, 29, 32, 64, 65] and found that IGH, IGK, and IGL amplification rates did not exceed 87% and were as low as 45% (Fig. 5A). Amplification rates for TCRA and TCRB were similar, ranging from 53% to 73% (Fig. 5A). The Ig primer set designed earliest by Sundling et al in 2012 [28] had the highest recovery of IGH sequences (76%), while the most recent Ig primer set designed by Rosenfeld et al in 2019 [32] had the highest recovery of IGK (87%) and IGL (71%). The TCRB specific primers designed by Li et al [65] recovered 73% of the TCRB sequences, while Chen et al [64] recovered 53% of TCRA sequences.

**Fig 5.**
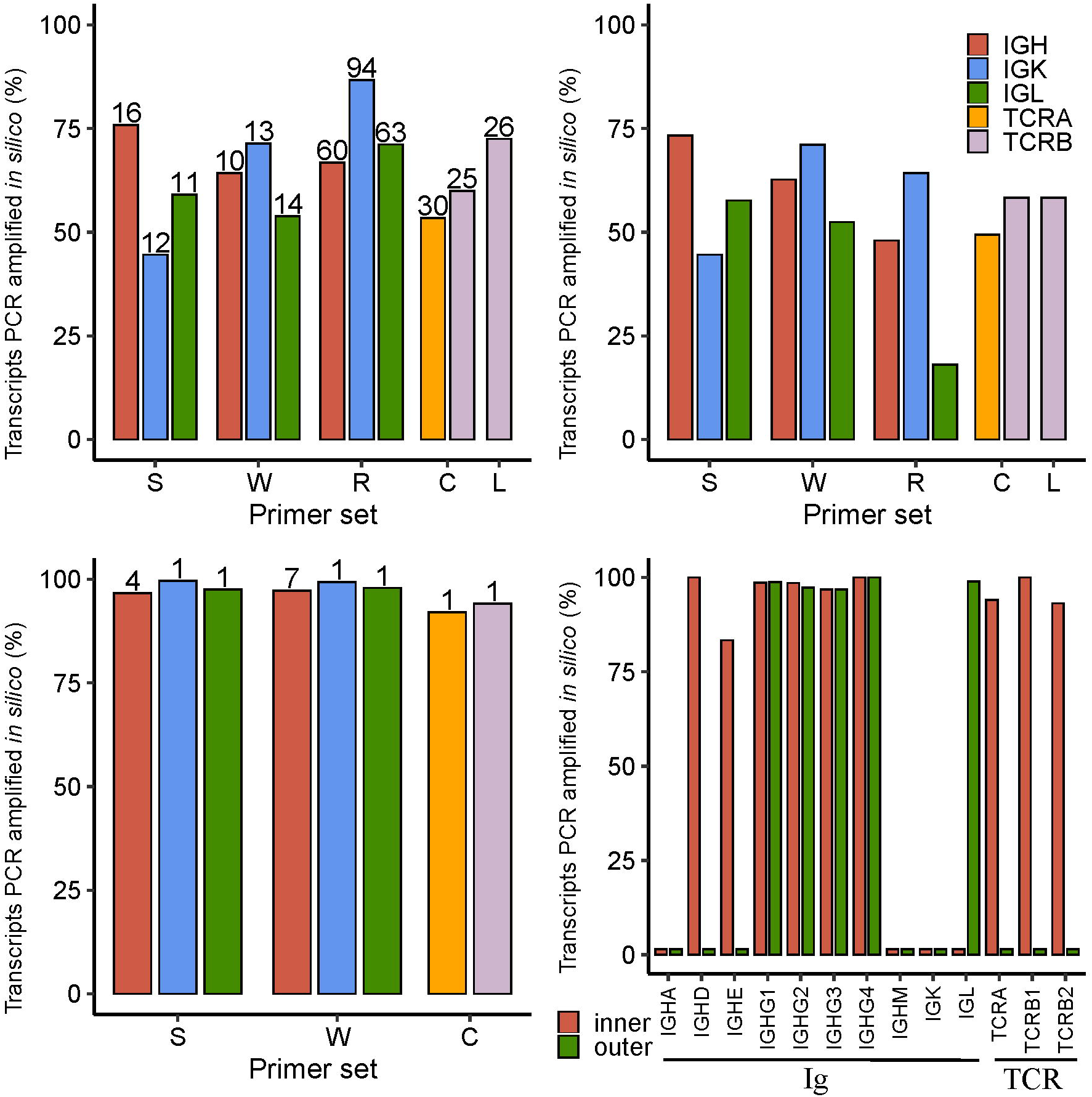
*In silico* PCR analysis of rhesus-specific primers from (S)undling et al [28], (W)iehe et al [29], (R)osenfeld et al [32], (C)hen et al [64], and (L)i et al [65] as well as human 10x B and T cell V(D)J reverse primers [66]. Only the inner primers were tested for S, W, while both inner and outer 10x primers were tested. The primers from C target both TCRA and TCRB, while those from L only target TCRB, as indicated by the bars shown in **A-C**. The percentage of transcripts amplified *in silico* are shown as bars for different primer sets and types of transcripts. **A)** Amplification where only primers targeting the V gene segment within the variable region from S, W, R, C, and L are tested with the total number of primers (after enumerating primers with degenerate nucleotides) shown above the bars. **B)** Amplification using forward (V gene) and reverse (J gene segment or constant region) primers from S, W, R, C, and L. **C)** Amplification where only primers targeting the constant region from S, W, and C are tested with the total number of primers (after enumerat ing primers with degenerate nucleotides) shown above the bars. R and L are not shown here as their reverse primers target J gene segments. **D)** Analysis of 10x B and T cell V(D)J inner and outer reverse primers. The different groups of transcripts (Ig and TCR) are noted below the bar plot.

As expected, when we assessed both forward and reverse primers together, the low percentages of sequences amplified largely remained the same (Fig. 5B). Notably, the percentages for Rosenfeld et al [32] and the percentage for Li et al [65] were even lower, as their reverse primers targeted J gene segments instead of the constant region. Indeed, when we tested the constant region primers alone (Fig. 5C), we found that Sundling et al [28], Wiehe et al [29], and Chen et al [64] had near perfect amplification rates, which was expected.

### Design of rhesus-specific B- and T-cell V(D)J assays for single cell analysis

Both our consensus profiling (Fig. 4) and *in silico* PCR analysis (Fig. 5A-C) showed that primers targeting variable regions are the source of bias in targeted amplification of rhesus Ig and TCR repertoires. Other methods, particularly RACE-seq and those using scRNA-seq, avoid this problem by only using primers targeting the constant region. Since there are no known single cell repertoire analysis assays for rhesus macaque, we first checked the efficiency of human B and T cell V(D)J assays developed by 10x Genomics [66] against our collection of Ig and TCR transcript sequences. We found that neither assay was able to fully capture this set of rhesus Ig and TCR transcript sequences (Fig. 5D). The human inner primers successfully amplified IGHD (100%), IGHE (83%), IGHG1 (99%), IGHG2 (98%), IGHG3 (97%), IGHG4 (100%), TCRA (94%), TCRB1 (100%), and TCRB2 (93%). Meanwhile, the outer primers only amplified IGHG1 (99%), IGHG2 (97%), IGHG3 (97%), IGHG4 (100%) and IGL (99%). Notably, TCRD, TCRG1, and TCRG2 were not included in this analysis, as 10x Genomics assays do not cover these transcripts.

We further assessed sequence homology between human and rhesus reference sequences for the constant regions. For each Ig and TCR isotype, we aligned our reference cDNA coding sequences for rhesus macaque with those available for human and evaluated the compatibility of their respective 10x human primers where relevant. For example, we aligned human IGHA1 and IGHA2 to the five rhesus IGHA consensus sequences (3 secreted, 2 membrane-bound) and examined the regions targeted by 10x human primers (Fig. 6A). Overall, the two human cDNA coding sequences had high similarity to the rhesus consensus sequences, averaging 89.7% identity. However, multiple nucleotide differences existed in the regions of rhesus reference sequences targeted by the 10x human outer and inner primers (Fig. 6A-C). These primer inconsistencies were not specific to IGHA, as all other human primers with the exception of the IGHG outer primer had mismatches with at least one of its corresponding rhesus reference sequences (data not shown). These results clearly indicated rhesus-specific primers are necessary to perform comparable single cell analysis for rhesus macaques.

**Fig 6.**
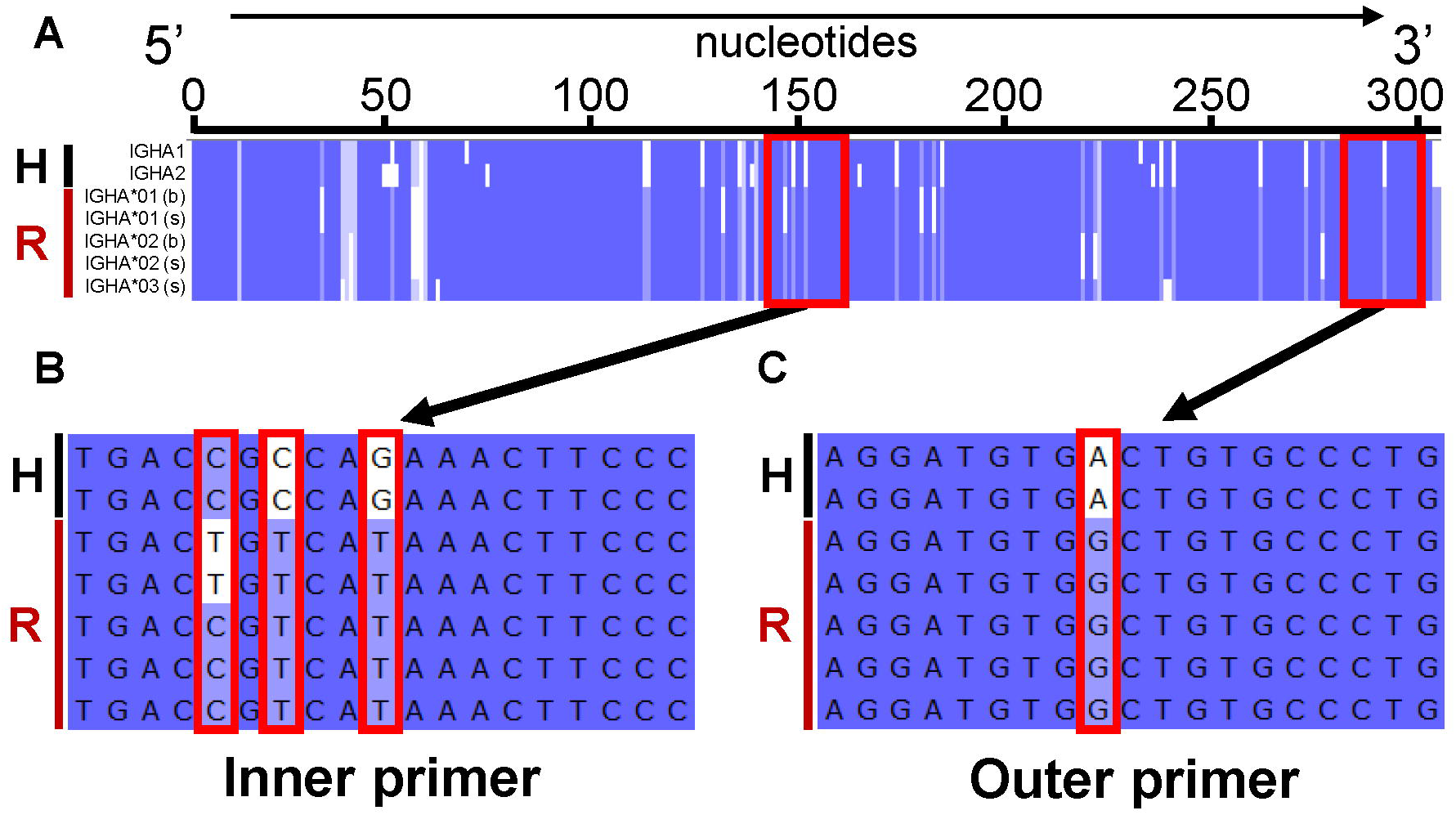
Comparison of human and rhesus IGHA reference sequences and illustration of human 10x primer target regions. A black H and red R, respectively highlight human and rhesus sequences on the left side of each panel. **A)** Multiple sequence alignment of putative human IGHA1 (J00220) and IGHA2 (J00221) CDSs with rhesus consensus cDNA coding sequences recovered (IGHA*01, IGHA*02, and IGHA*03) in either secreted (s) or membrane-bound (b) form. The target sites of human 10x primers are denoted by red boxes. **B-C)** Zoom in of the inner and outer primer target sites, where mismatches between the primer and rhesus consensus sequence(s) are denoted by red boxes.

To properly design rhesus-specific Ig and TCR constant region primers that are compatible with respective 10x human V(D)J assay designs, we first selected existing primers that were free of mismatches, which included the 10x human IGHG outer primer and rhesus primers designed by Sundling et al [28] (Supplementary data). Notably, some primers that successfully *in silico* PCR amplified Ig/TCR transcripts contained one or more mismatches thus requiring updated designs (Materials and Methods). For isotypes that lacked an inner and/or outer primer, we designed new primers with melting temperatures (Tms) compatible with that of the 10x forward primer (Supplementary data). Similar to the Ig primers, 10x human TCR primers were also largely incompatible with the rhesus consensus sequences. We found that only the inner primer for TCRB had a perfect match and thus designed all other primers for rhesus macaque (Supplementary data). We also included primers that target TCRD and TCRG in the final set of our rhesus-specific T-cell V(D)J primers, which are not included in the human 10x assays [66].

To ensure these newly designed primers can amplify rhesus transcripts experimentally, for each of them we designed a corresponding forward primer that was upstream but still within the constant region (Supplementary data). Then we tested all these custom inner and outer primers by pairing with their forward primers via PCR (Materials and Methods). We obtained the desired product size for each assay (Supplementary Fig. 3), confirming that these rhesus specific primers newly designed for single cell based V(D)J assays targeted rhesus Ig/TCR transcripts as expected. In some assays, we also observed additional bands, but the desired bands tended to be visibly the most promine nt (7 of 8 outer primers and 2 of 5 inner primers tested). Many of those additional bands were larger than the desired product size (Supplementary Fig. 3), suggesting some forward primers might have shared certain sequence similarities with the highly diverse upstream variable regions. The actual single cell assays use nested template switch PCR; therefore, these forward primers are not needed and such off-target amplification would not be a concern. Together, these results indicate that the primers we designed will be useful for single cell assays and compatible with the 10x Genomics platform.

## Discussion

In this study, we profiled Ig and TCR repertoires using long read transcript sequencing of tissues collected from Indian-origin rhesus macaques, one of the most widely used NHP models. Since we obtained high-quality, full-length sequences for individual Ig and TCR transcripts, no sequence assembly was necessary. This unbiased profiling analysis allowed us to compile the first complete reference set for the constant regions of all expected rhesus Ig and TCR isotypes and chain types based on homology with human, including three newly identified reference sequences. While most of publicly available rhesus Ig and TCR constant region sequences were computationally predicted from genomic sequences and/or partial sequences, our reference sequences were from directly sequenced transcripts and full length. We were able to define the coding regions for these constant regions, yielding full coding sequences as well as cDNA sequences containing 3’ UTRs.

An immediate benefit of having this complete constant region reference is improved accuracy of rhesus Ig/TCR transcripts identification. For example, we found that about 39% of rhesus TCR transcripts that we initially identified by the commonly used combination of IgBLAST and IMGT databases were actually Ig transcripts, based on the alignment of their constant region portion against these reference sequences. This miss-assignment could be due to several reasons. First, the current IMGT database has a limited number of TCR variable region germline sequences for rhesus macaque. Second, the alignment of rhesus Ig/TCR transcripts to the human variable region reference is not sufficiently reliable. The incomplete 5’ end of some rhesus Ig/TCR transcript sequences was also a contributing factor. In the future, we expect that our reference sequences will improve identification of novel germline variable genes extracted from recombined Ig and TCR sequences in rhesus macaque, as the number of false positives will be substantially abrogated.

These full-length Ig/TCR transcript sequences offered insights into the rhesus immune system. For the Ig constant regions, we identified both cell membrane-bound and secreted forms that we differentiated by distinct 3’ end splicing. This demonstrates our ability to detect the alternative splicing events among these multi-exonic regions. As the genomic assembly and annotation of these constant region loci improves, long read sequencing may enable discovery of similar alternative exon usage as described for human Ig heavy chain constant region genes [71]. In addition, we recovered three different allotypes for IGHA with significant variation in the hinge region. When this hinge region is longer, the avidity for antigen interactions is increased at the expense of increased susceptibility to pathogen-derived proteases known to target the hinge region [72]. We suspect that our ability to identify hinge region variation was driven by the high heterogeneity of the alpha constant region [69] as well as its highest frequency among the Ig isotypes we recovered. Given the interspecies variability of IGHG between macaques and humans [73] and the heterogeneity of rhesus IGHC genes in general [16], it is imperative that future rhesus Ig gene analyses also consider this allotypic diversity to improve the translatability of NHP models.

Since we did not use any targeted experimental approaches to obtain these full-length Ig/TCR transcripts, we were able to examine the diversities of rhesus Ig/TCR variable regions in an unbiased manner. We found that large diversities existed across the entire variable regions of rhesus Ig and TCR transcripts. This is especially relevant, since available assays for profiling rhesus Ig/TCR repertoires have relied on the consensus primers derived from a limited number of available rhesus variable region sequences. Previous benchmarking analyses for targeted repertoire sequencing in human used either spiked-in samples [43] or samples of unknown composition [42], motivating us to leverage our collected sequences for a truly unbiased assessment of conventional rhesus-specific MPCR designs for Ig [28, 29, 32] and TCR [64, 65]. MPCR targets specific V gene segments (forward primer) and either specific J gene segments or constant regions (reverse primer) to generate amplicon libraries. Not surprisingly, we found that all of these assays failed to cover a significant percentage of these rhesus Ig/TCR transcripts. We do not contend that this method be avoided altogether moving forward, but additiona l improvements will be necessary. For example, databases of variable region gene segments can be improved using the IgDiscover tool [18]. Targeting the more conserved 5’ UTR of V genes may reduce the amount of primers needed in MPCR designs and yield amplico ns covering the entire variable region but still need to be designed for rhesus macaque [34]. New algorithms such as the recently described version of the tool for Ig Genotype Elucidation via Rep-Seq (TIgGER) for identifying human Ig [74] can also improve germline allele assignment if adapted for rhesus macaque.

Alternatively, scRNA-seq and RACE-seq only use primers that target the constant regions of these Ig/TCR transcripts. In particular, scRNA-seq combines repertoire analysis and transcriptome profiling in single B and T cells [51–54], and it can identify chain pairs within single cells [75, 76]. Our analysis showed that human-specific constant region primers from commercially available assays were not compatible for use in rhesus macaque. For these reasons, we elected to design the first single cell based B- and T-cell V(D)J assays for rhesus macaque. We leveraged the complete set of rhesus constant region reference sequences compiled here to ensure a broad coverage of rhesus Ig/TCR diversity and experimentally validated the newly designed rhesus-specific primers. Our assays are fully compatible with the standard protocols from 10x Genomics, currently a popular platform for single cell analysis. These primers can also be used or modified for RACE-seq applications, considering similar library preparation techniques.

More broadly, this study also demonstrated an accessible strategy for developing immune repertoire resources and assays for less studied organisms. Traditionally, the development of Ig/TCR resources would start with collecting large numbers of species-specific transcript and germline sequences. Since Ig/TCR loci are among the most complex regions in the genome due to their duplicated and polymorphic nature, proper sequencing and assembly of these genomic regions requires sophisticated efforts [77]. Consequently, complete assemblies are only available for selected model organisms such as human and mouse; meanwhile, these loci still do not have complete assemblies for rhesus macaque despite recent efforts [16]. The PacBio Iso-Seq approach we used here sequences full-length Ig/TCR transcripts directly and in a high-throughput manner, bypassing transcript assembly issues. The performance of this strategy could be further improved as well. For example, compared to the whole tissue sample used here, use of isolated B and T cells may significantly improve the recovery of these repertoire sequences. The custom computational solution we developed and describe here can be modified for other species by replacing constant region references from a related species to identify putative species-specific Ig/TCR transcript sequences. However, additional information like Ig/TCR germline allele assignment, the genomic order of individual genes, and copy number variations will still require complete genomic sequencing and assembly.

In this study, we systematically profiled the diversity of rhesus Ig and TCR genes using full-length transcriptome sequencing. We performed benchmark analysis of common MPCR based targeted sequencing strategies for rhesus macaque and demonstrated that available MPCR assays do not cover the rhesus repertoire diversity adequately. Our construction of a complete reference set for rhesus macaque Ig/TCR constant regions enabled the design of the first rhesus-specific single cell based V(D)J assays, addressing many of the current technical limitations of rhesus macaque repertoire analysis. Furthermore, our approach circumvents the computational challenges concomitant with the genomic assembly of these complex immune genes by combining long read sequencing and new computational solutions. This study thus represents a new direction for developing nascent immune resources.

## Abbreviations

CCS: circular consensus sequence
NHP: non-human primate
Ig: immunoglobulin
IGH: Ig heavy chain
IGHA: IGH alpha
IGHD: IGH delta
IGHE: IGH epsilon
IGHG: IGH gamma
IGHM: IGH mu
IGK: Ig kappa chain
IGL: Ig lambda chain
TCR: T cell receptor
TCRA: TCR alpha
TCRB: TCR beta
TCRD: TCR delta
TCRG: TCR gamma
IMGT: the international ImMunoGeneTics information system
MPCR: multiplex PCR
RACE: rapid amplification of cDNA ends

