## Supplementary Figures for "Systematic profiling of full-length immunoglobulin and T-cell receptor repertoire diversity in rhesus macaque through long read transcriptome sequencing"

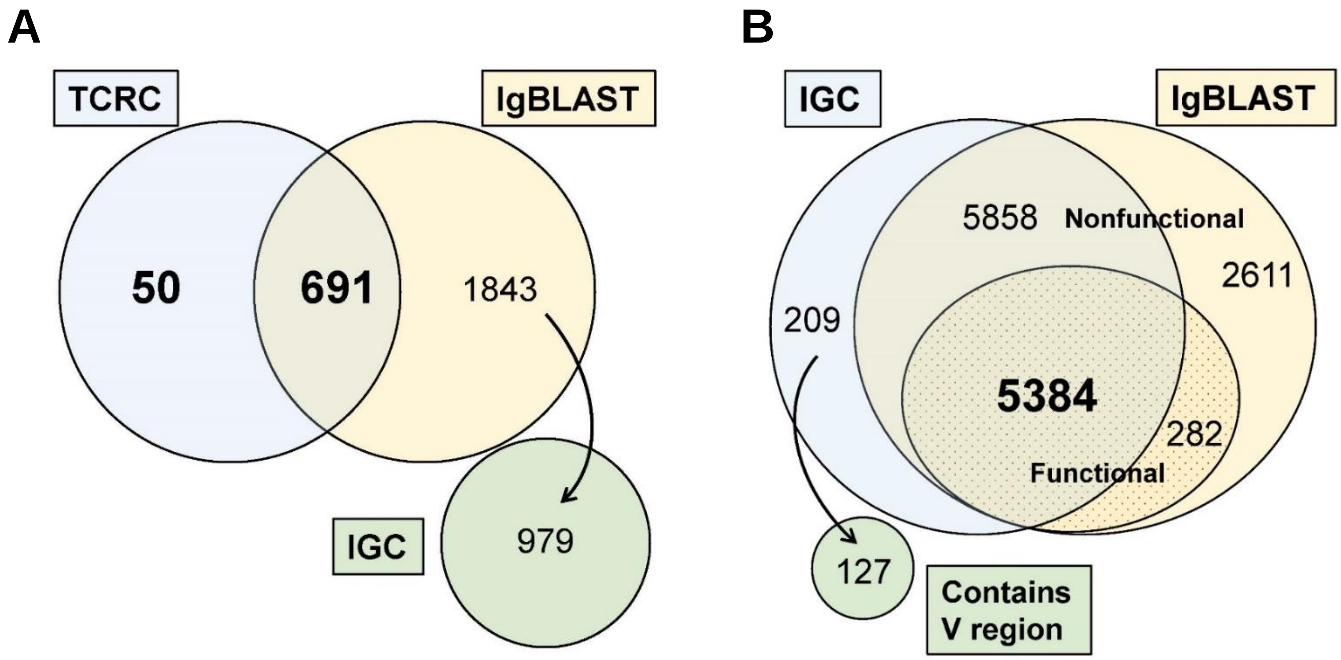


**Supplementary Figure 1.** Venn diagrams depicting comparisons of how CCS reads assigned to Ig/TCR depending on the method used. The yellow circles represent reads assigned to Ig/TCR using the IgBLAST tool, which makes assignments based on mapping the variable region to a database of germline reference sequences of known V, D, and J gene segments. Blue circles indicate reads aligned to our custom reference of IG/TCR constant regions. **A)** Assignment of reads to TCR using the IgBLAST tool with human IMGT reference sequences. Sequences unique to the IgBLAST search were also mapped to the IGC database to reveal those that were in fact Ig sequences (green circle). **B)** Assignment of reads to Ig using the IgBLAST tool with rhesus IMGT reference sequences. These reads were considered functional (plain yellow) or nonfunctional (dotted yellow), which is decided based on the presence or absence of premature stop codons and if the read is considered to be in frame with the reference germline sequences it mapped to. Sequences unique to the IGC database search were further analyzed to reveal sequences with sufficiently long variable regions (> 100nt) for detection by IgBLAST (green circle).

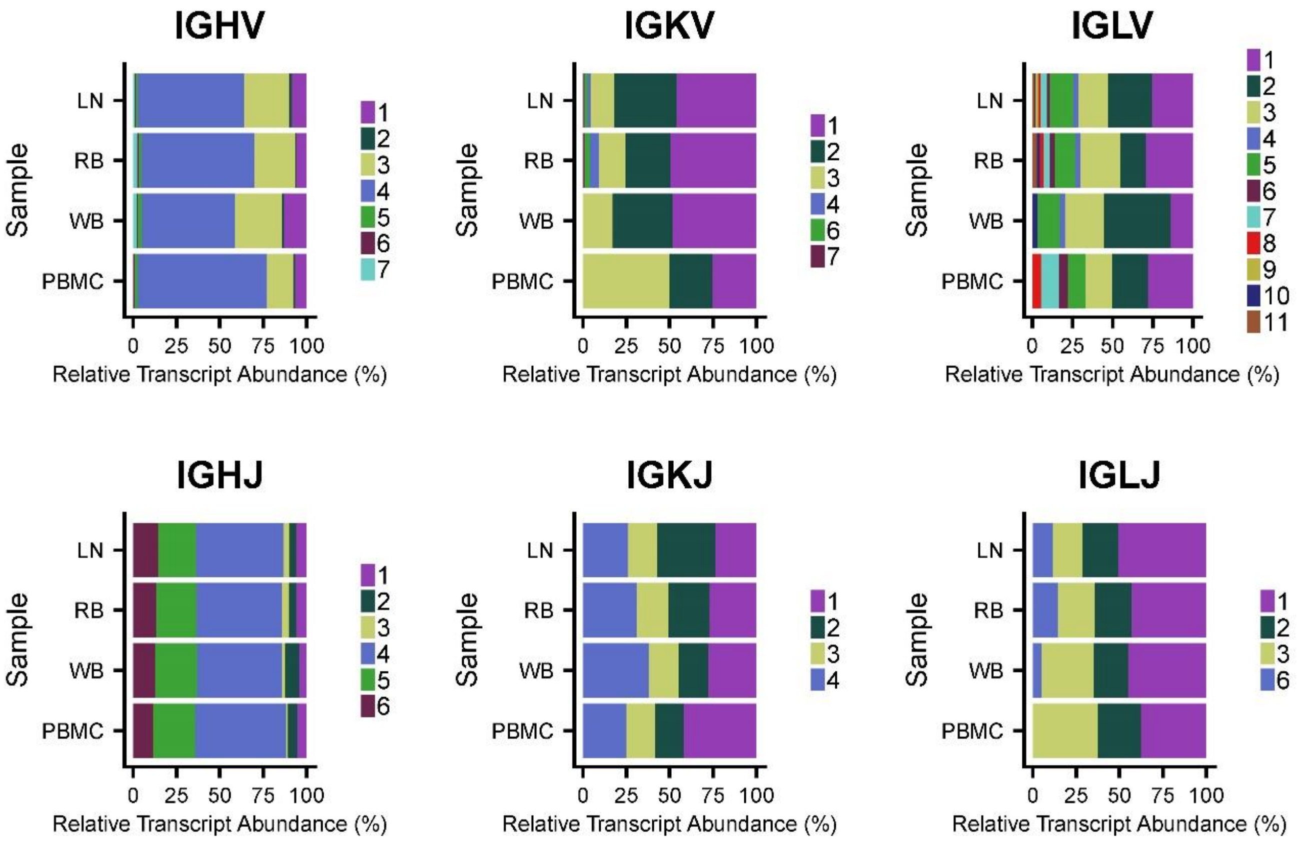


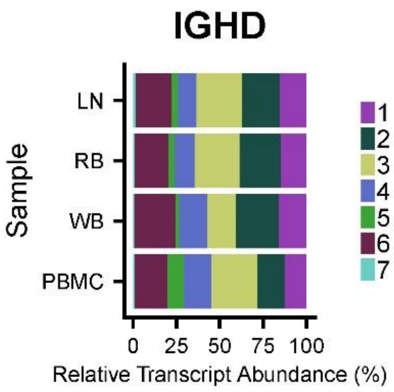


**Supplementary Figure 2.** Relative transcript abundances of the various Ig V, D and J gene segments. In each stack bar plot, Samples are displayed vertically and the various gene families for the type of gene segment are colored differently as described in the legend to the right of the plot. Gene families (e.g. IGHV1) are clusters of the corresponding gene segment type (IGHV) that have high sequence similarity.

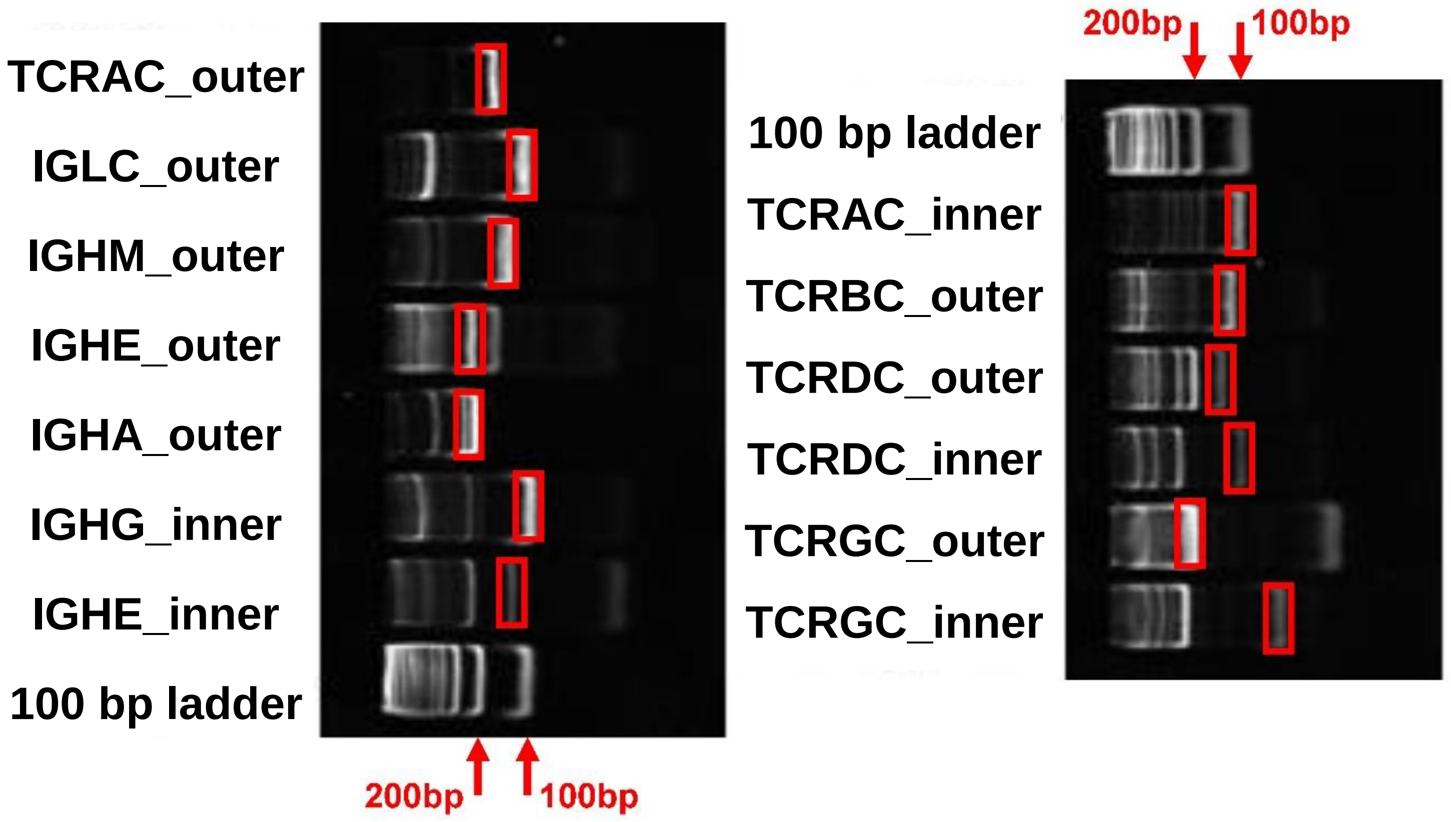


**Supplementary Figure 3.** Polyacrylamide gel images show the PCR products from lymph node RNA using primers specific to rhesus Ig and TCR constant regions sequences. The desired bands are outlined by red boxes, with gene rulers showing the approximate band sizes (bottom of left gel image, top of right gel image). Since all desired bands were detected, the other tissues used in initial PacBio sequencing were not tested. The reverse primers are designed for use in a nested template switch PCR assay, while the forward primers are designed solely for the purposes of this assay. Lane names indicate the constant region targeted (e.g. TCRAC) and whether it is the inner or outer primer being tested.

| Tissue | Type of CCS read | Number of CCS reads | % Ig transcripts | | % TCR transcripts | |
|---|---|---|---|---|---|---|
| | | | IgBLAST | IgBLAST + F + IGC | IgBLAST | TCRC |
| Lymph Node | Full | 234,348 | 2.311 | 2.029 | 0.432 | 0.160 |
| | Partial | 503,492 | | | | |
| Peripheral Blood Mononuclear Cells | Full | 183,174 | 0.291 | 0.154 | 0.254 | 0.130 |
| | Partial | 401,001 | | | | |
| Rectal Biopsy | Full | 261,011 | 2.492 | 2.028 | 0.254 | 0.008 |
| | Partial | 535,317 | | | | |
| Whole Blood | Full | 243,466 | 0.273 | 0.180 | 0.162 | 0.044 |
| | Partial | 442,883 | | | | |
| Total | Full | 921,999 | 1.423 | 1.168 | 0.275 | 0.080 |
| | Partial | 1,882,693 | | | | |

**Supplementary Table 1.** PacBio long read based full-length transcriptome sequencing yields Immunoglobulin (Ig) and T cell receptor (TCR) sequences. Four tissues were sequenced and CCS reads were generated from unique sequencing molecules. CCS reads are classified as full if they contain the 5’ primer, 3’ primer, and polyA tail, while partial sequences lack one of these components. The percentages of CCS reads identified as Ig and TCR sequences are shown on the right side of the table. For Ig, we show the raw IgBLAST counts using rhesus and human annotation (IgBLAST) as well as the final counts of functional IgBLAST hits that also aligned to our custom Ig constant region database (IgBLAST + F + IGC). For TCR, we show the raw IgBLAST counts using human annotation (IgBLAST) as well as the final counts of transcripts that aligned to our custom TCR constant region database (TCRC).
